# Limitation of dogwhelk consumption of mussels by crab cues depends on dogwhelk density and cue type

**DOI:** 10.1101/123653

**Authors:** Melanie L. Boudreau, Ricardo A. Scrosati, Melisa C. Wong

## Abstract

Predator nonconsumptive effects (NCEs) on prey activity are common in nature. Upon sensing predator cues, a common prey response is to reduce feeding to avoid being detected by predators. Using an aquatic system, this study investigated how prey density and predator cue type affect predator NCEs on prey feeding. Prey density was investigated because, as it increases, the individual risk of being preyed upon decreases, which may reduce NCEs if prey can detect conspecifics. Predator cue type was investigated because waterborne cues would trigger weaker NCEs than waterborne and tactile cues combined, as predation risk may be perceived by prey to be stronger in the second case. Specifically, a factorial experiment tested the hypotheses that (i) increasing dogwhelk (prey) density reduces the limitation that crab (predator) chemical cues can have on dogwhelk consumption of mussels and that (ii) chemical and tactile crab cues combined limit dogwhelk feeding more strongly than chemical crab cues alone. The results broadly supported these hypotheses. On the one hand, crab chemical cues limited the per-capita consumption of mussels by dogwhelks at a low dogwhelk density, but such NCEs disappeared at intermediate and high dogwhelk densities. On the other hand, the combination of chemical and tactile cues from crabs caused stronger NCEs, as dogwhelk consumption of mussels was negatively affected at all three dogwhelk densities. The structurally complex mussel beds may provide not only food for dogwhelks but a refuge from crab predation that allows dogwhelk density to limit crab NCEs when mediated by waterborne cues. Overall, this study suggests that prey evaluate conspecific density when assessing predation risk and that the type of cues prey are exposed to can affect their interpretation of risk.

## Introduction

Predators regulate prey populations through direct consumption, but they also often have nonconsumptive effects (NCEs). Upon detection of predator cues, prey commonly react by moving away or reducing feeding activities to reduce predation risk (Keppel and Scrosati, 2004; Molis et al., 2011; Hossie et al., 2017). Such responses may, in turn, favor species at a lower trophic level as consumption by the intermediate level decreases due to the NCEs from the top predator. As predator NCEs can influence many prey organisms simultaneously, the cascading effects on communities can be extensive (Preisser et al., 2005; Madin et al., 2016). Thus, understanding the factors that affect the occurrence of NCEs is a central theme in NCE research (Weissburg et al., 2014).

A number of studies have found that conspecific prey density may influence the occurrence of predator NCEs on prey (Ferrari et al., 2010; Guariento et al., 2015). For example, on marine shores, the presence of adult barnacles or a high density of barnacle recruits neutralize the limitation that cues from predatory dogwhelks would otherwise exert on barnacle recruitment (Ellrich et al., 2015, 2016). The absence of such NCEs in the presence of barnacle conspecifics likely occurs because pelagic barnacle larvae seeking settlement are attracted by chemical cues from conspecific recruits and adults (Crisp and Meadows, 1962; Matsumura et al., 2000). Benthic conspecifics would indicate to settling larvae that local conditions are adequate for survival and development (Clare, 2011). Prey larvae thus seem to assess conspecific density as part of their evaluation of future predation risk as settled larvae develop into adults. Microcosm experiments with other aquatic species have found that predator NCEs on prey activity and growth also weaken with prey density (Turner, 2004; Van Buskirk et al., 2011). The importance of prey density for the occurrence of predator NCEs has also been recognized from a theoretical viewpoint (Peacor, 2003).

In contrast to those studies, however, other studies have found that increasing prey density does not eliminate predator NCEs, raising the question of under what circumstances does prey density matter. These other studies used green crabs (*Carcinus maenas*), dogwhelks (*Nucella lapillus*), and barnacles (*Semibalanus balanoides*). Green crabs consume mussels (*Mytilus edulis*) and also dogwhelks (Ropes, 1968; Elner, 1978; Hughes and Elner, 1979), while dogwhelks consume barnacles and mussels (Dunkin and Hughes, 1984; Hughes and Dunkin, 1984; Crothers, 1985). Dogwhelks can detect chemical cues released by crabs fed either mussels or dogwhelks (Large and Smee, 2010) and also metabolites released by conspecific dogwhelks while they consume mussels (Hughes and Dunkin, 1984; Large and Smee, 2010). Experimental work has shown that chemical cues from green crabs reduce the per-capita consumption of barnacles by dogwhelks, but doubling or even tripling dogwhelk density does not neutralize such NCEs (Trussell et al., 2003, 2006). Later work showed that the limitation of dogwhelk consumption caused by crab cues is stronger when dogwhelks feed on barnacles than when they feed on mussels (Trussell et al., 2008). It was suggested that, because mussel beds are structurally more complex than the relatively flat barnacle stands, dogwhelks would find better refuge opportunities in mussel beds, prompting dogwhelks to react less strongly to waterborne crab cues. Therefore, an increasing dogwhelk density might reduce crab NCEs on dogwhelk feeding when the dogwhelks consume mussels from extensive beds. This paper tests this hypothesis experimentally using the species mentioned above. Although a recent study suggested that refuge availability may intensify predator NCEs because prey in refuges often have lower access to food (Orrock et al., 2013), mussel beds provide both a refuge as well as food (in contrast to inert refuges; see also Donelan et al., 2017), which supports testing the above hypothesis.

In addition to prey density, cue type may also affect the occurrence of NCEs (Stauffer and Semlitsch, 1993; Chivers et al., 2001; Luttbeg and Trussell, 2013). For example, predator chemical cues alone may indicate to prey a less immediate risk of predation than the combination of chemical and tactile (predators touching the prey) cues from the predators. Therefore, this paper also evaluates whether crab cue type interacts with dogwhelk density to influence crab NCEs. Thus, the second hypothesis of this study is that dogwhelk density weakens crab NCEs on dogwhelk feeding more strongly under chemical crab cues alone than under chemical and tactile crab cues combined.

## Materials and methods

The hypotheses were tested through a laboratory experiment conducted between late summer and early fall. The experimental units that contained the crabs, dogwhelks, and mussels were glass aquaria of 54 L (60 cm × 30 cm × 30 cm) with flow-through seawater running at a rate of 2 L min^-1^. The photoperiod was 12:12 and seawater temperature averaged 12.5 °C. All organisms were collected on the Atlantic coast of Nova Scotia, Canada. The mussels (15–20 mm long) were collected at the rocky intertidal zone of Chebucto Head (N44 40.967, W63 36.790), the dogwhelks (18–23 mm long) at Blandford (N44 28.666, W64 5.897), and the green crabs (50–60 mm of carapace width) at Little Port Joli Lagoon (N43 52.315, W64 49.381). These species coexist along this shore, but doing the collections at these locations facilitated obtaining enough organisms for the study. The size ranges were selected based on preliminary trials that identified appropriate mussel sizes to maximize dogwhelk feeding and to ensure crab predation on mussels in the treatment with chemical and tactile crab cues (described below). For consistency, only male crabs without missing limbs and dogwhelks and mussels with intact shells were used. Once collected in the field, the organisms were kept in laboratory tanks with flow-through seawater for 12 days before the start of each of the three experimental blocks described below. During that acclimation periods in the tanks, the crabs were fed a combination of mussels and whitefish, while the dogwhelks were fed mussels. Crabs and dogwhelks were subjected to a starvation period of five days before the start of each experimental block to standardize their hunger level.

The experiment evaluated the effects of dogwhelk density and crab cue type following a randomized complete block design with replicated treatments within blocks (Quinn and Keough, 2002). Dogwhelk density included three treatments: low (6 dogwhelks per aquarium), intermediate (11 dogwhelks), and high density (17 dogwhelks), corresponding to 33, 61, and 94 dogwhelks m^-2^. These densities are within the natural range found on the coast where the organisms were collected. Crab cue type included three treatments: no cues (NC), chemical cues (CC), and tactile and chemical cues (TCC). The NC treatment represented a crab absence in an aquarium. The CC treatment represented the occurrence of a crab in a perforated circular container (15.7 cm in diameter and 7.6 cm tall) in an aquarium, enabling the crab's chemical cues to reach the dogwhelks without allowing the crab to touch the dogwhelks. The TCC treatment represented a free-living crab in an aquarium, the crab being able to touch the dogwhelks. Each replicate aquarium contained 400 mussels, simulating the extensive mussel patches that are common in the habitats where the organisms were collected (Arribas et al., 2014). This number of mussels also ensured that they did not become limiting (less than 200 mussels per aquarium) during the experiment, as found by preliminary trials. Each crab in the CC treatment was fed six dogwhelks (placed in the circular container at the beginning of the experiment), while the crabs in the TCC treatment were able to feed on the mussels that were also available for the dogwhelks (the crabs in this treatment did not eat dogwhelks). The experiment used three blocks, each one lasting for seven days and consisting of three independent replicates of each of the nine treatments described above (three levels of dogwhelk density crossed with three levels of crab cue type), yielding a total of nine replicates for each treatment for the experiment. Separate aquaria not used for the experiment included 400 mussels and a free-living crab but no dogwhelks, which confirmed that the presence of dogwhelks did not affect the consumption rate of mussels by crabs. Three other aquaria contained each 400 mussels in the absence of dogwhelks and crabs to quantify the appearance of empty shells unrelated to predation.

At the end of each weekly block, the per-capita rate of consumption of mussels by dogwhelks was calculated for each aquarium by observing the condition of all mussels. A dogwhelk commonly drills a borehole through the shell of a mussel to consume its internal tissues (Carriker and Williams, 1978). Less often, a dogwhelk can consume a mussel by forcing its proboscis between the mussel valves, without leaving a borehole (M. L. Boudreau, pers. obs.; Rovero et al., 1999). At the end of each weekly block, the mussels from every aquarium were sorted into nine categories: (1) alive, (2) empty shell with no borehole (indicating either natural mortality or full dogwhelk consumption between the mussel valves), (3) empty shell with one borehole, (4) empty shell with two boreholes, (5) partial internal remains with no borehole, (6) partial internal remains with one borehole, (7) partial internal remains with two boreholes, (8) fragmented shell with boreholes (indicating a combined crab and dogwhelk consumption in the TCC treatment), and (9) gaping mussel with all internal biomass and no borehole (indicating natural mortality). Fragmented shells with no boreholes were found only in the TCC treatment and suggested consumption of mussels only by crabs, so the number of such shells was not used to calculate dogwhelk consumption rates.

The per-capita rate of consumption of mussels by dogwhelks (mussels dogwhelk^-1^ week^-1^) was calculated for each aquarium using this formula: {[(*N*_2_ − *N*_d_) + *N*_3_ + *N*_4_ + (*N*_5_ / 2) + (*N*_6_ / 2) + (*N*_7_ / 2) + (*N*_8_ × 0.25)] / *D*}. The expressions *N*_2_ to *N*_8_ refer to the number of mussels found respectively for categories 2 to 8 described above, *N*_d_ refers to the average number of mussels that died naturally leaving empty shells in the three aquaria without dogwhelks or crabs, and *D* refers to the number of dogwhelks. The formula subtracts *N*_d_ from *N*_2_ to determine as realistically as possible the number of category-2 mussels that were consumed by dogwhelks. Even though category-4 mussels had two boreholes, their number (*N*_4_) was not divided by 2 because the dogwhelk per-capita consumption rate must necessarily be calculated, when using data for fully consumed mussels, by dividing the number of such mussels by the number of dogwhelks in the aquarium. As mussels from categories 5, 6, and 7 were partially consumed, an average of 50 *%* of their internal biomass was estimated to remain, so their respective numbers (*N*_5_, *N*_6_, and *N*_7_) were divided by 2. Even though category-7 mussels had two boreholes, their number (*N*_7_) was not further divided by 2 for the reason given above for *N*_4_. The number of category-8 mussels (*N*_8_) was multiplied by 0.25 because the mussels consumed by both a crab and a dogwhelk (in the TCC treatment) were mostly eaten (ca. 3/4) by the crab, which often interrupted dogwhelk feeding at an early stage.

Once the per-capita dogwhelk consumption rate of mussels was determined for each aquarium, the effects of dogwhelk density and crab cue type were analyzed through a factorial analysis of variance (ANOVA; Quinn and Keough, 2002). Dogwhelk density was considered as a fixed factor with three levels (low, medium, and high), crab cue type as a fixed factor with three levels (NC, CC, and TCC), and block as a random factor with three levels (the three weekly periods). The assumptions of normality and homoscedasticity were verified with the Kolmogorov-Smirnov test and Cochran’s *C*-test, respectively (Quinn and Keough, 2002). Because blocks and the interaction terms including blocks yielded *P* values higher than 0.2 (Table 1), those sources of variation were removed from the model and a final ANOVA was done without them (Winer et al., 1991). Although the interaction term was not significant in the final ANOVA, the observed trends in the data suggested an apparent dependence of crab cue effects on dogwhelk density. To examine that possibility in more detail, a post-hoc power analysis was conducted for the interaction term (Zar, 1999). Because power was low for the interaction (see Results), tests of simple effects were done to evaluate crab cue effects separately for each dogwhelk density. Each of such tests was done as a one-way ANOVA using the error term from the final factorial ANOVA, followed by Tukey HSD tests to compare crab cue treatments (Quinn and Keough, 2002). The data analyses were conducted with SYSTAT 12.

**Table 1.**
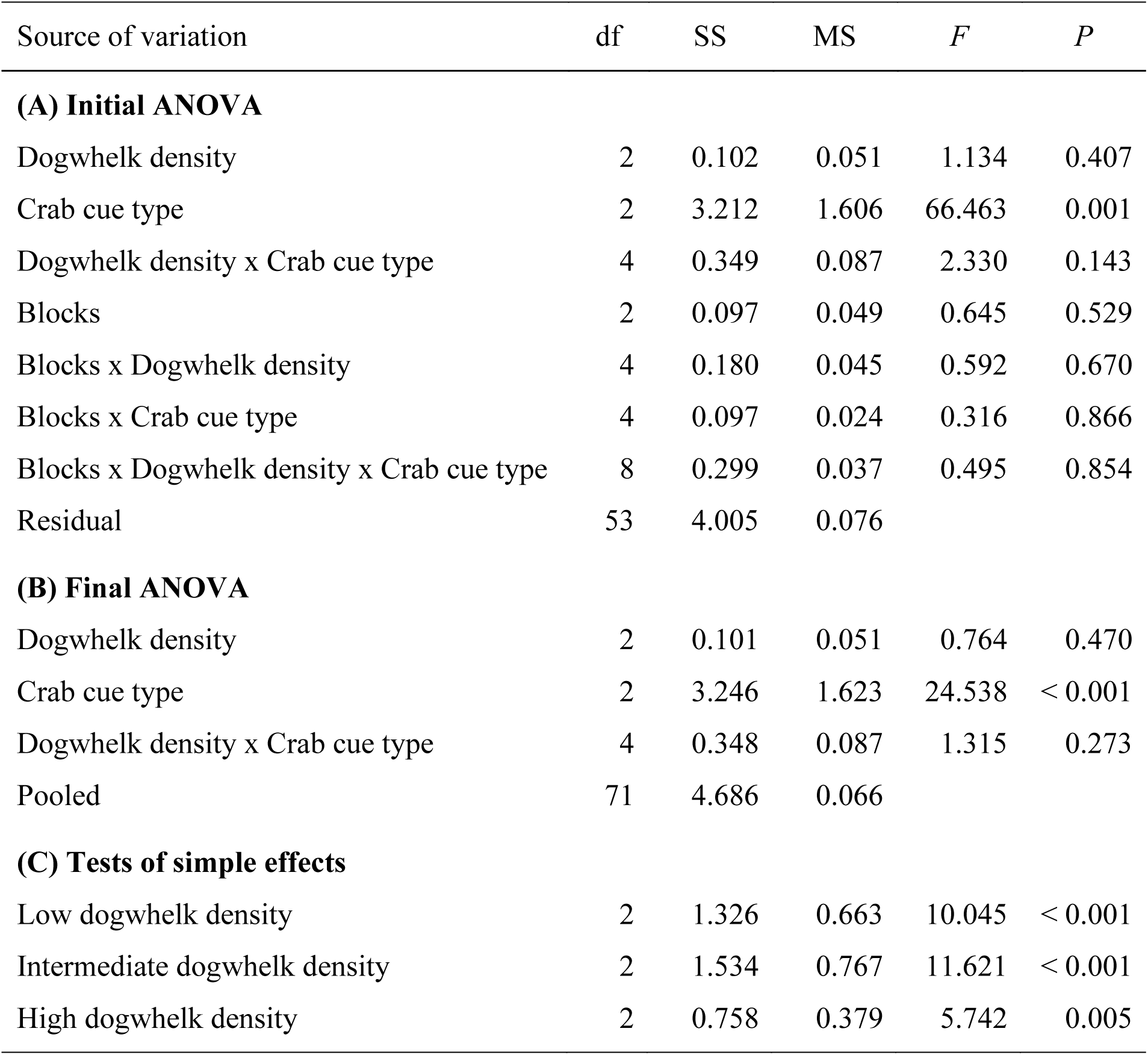
Summary results of (A) the initial ANOVA that tested the effects of dogwhelk density, crab cue type, and blocks on the per-capita consumption rate of mussels by dogwhelks, (B) the final ANOVA after the sum of squares for the sources of variation that included blocks was pooled with the residual variation (because their *P* > 0.2), and (C) the tests of simple effects done at the three evaluated levels of dogwhelk density. See the Results section for the rationale that supported performing the tests of simple effects.

| Source of variation | df | SS | MS | *F* | *P* |
| --- | --- | --- | --- | --- | --- |
| **(A) Initial ANOVA** |  |  |  |  |  |
| Dogwhelk density | 2 | 0.102 | 0.051 | 1.134 | 0.407 |
| Crab cue type | 2 | 3.212 | 1.606 | 66.463 | 0.001 |
| Dogwhelk density x Crab cue type | 4 | 0.349 | 0.087 | 2.330 | 0.143 |
| Blocks | 2 | 0.097 | 0.049 | 0.645 | 0.529 |
| Blocks x Dogwhelk density | 4 | 0.180 | 0.045 | 0.592 | 0.670 |
| Blocks x Crab cue type | 4 | 0.097 | 0.024 | 0.316 | 0.866 |
| Blocks x Dogwhelk density x Crab cue type | 8 | 0.299 | 0.037 | 0.495 | 0.854 |
| Residual | 53 | 4.005 | 0.076 |  |  |
| **(B) Final ANOVA** |  |  |  |  |  |
| Dogwhelk density | 2 | 0.101 | 0.051 | 0.764 | 0.470 |
| Crab cue type | 2 | 3.246 | 1.623 | 24.538 | < 0.001 |
| Dogwhelk density x Crab cue type | 4 | 0.348 | 0.087 | 1.315 | 0.273 |
| Pooled | 71 | 4.686 | 0.066 |  |  |
| **(C) Tests of simple effects** |  |  |  |  |  |
| Low dogwhelk density | 2 | 1.326 | 0.663 | 10.045 | < 0.001 |
| Intermediate dogwhelk density | 2 | 1.534 | 0.767 | 11.621 | < 0.001 |
| High dogwhelk density | 2 | 0.758 | 0.379 | 5.742 | 0.005 |

## Results

The final factorial ANOVA (Table 1) indicated that the type of crab cue significantly affected the per-capita rate of consumption of mussels by dogwhelks. Although the interaction between crab cue type and dogwhelk density was not significant, a post-hoc power analysis revealed that the statistical power associated to testing that interaction was lower than 0.25. The tests of simple effects that evaluated that interaction in more detail revealed that the influence of crab cue type depended on dogwhelk density. Crab chemical cues (CC) limited dogwhelk consumption rate at low dogwhelk density, but such an effect disappeared at intermediate and high dogwhelk densities (Table 1, Fig. 1). On the contrary, the combination of tactile and chemical cues from crabs (TCC) limited dogwhelk consumption rate at the three studied levels of dogwhelk density (Table 1, Fig. 1).

**Fig. 1.**
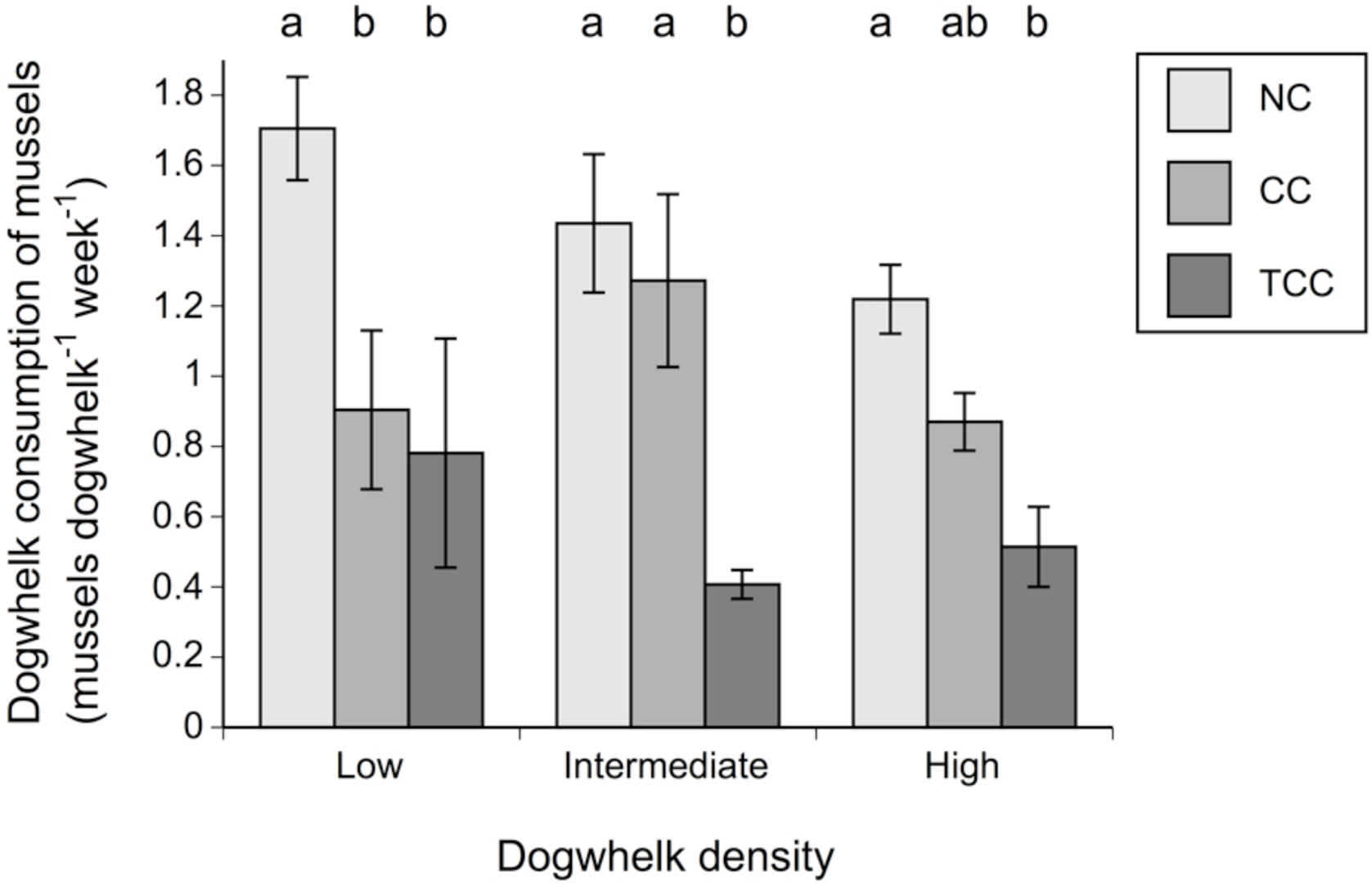
Per-capita rate of consumption of mussels by dogwhelks (mean ± SE) at the three levels of dogwhelk density and crab cue type considered in this study. For each level of dogwhelk density, significant differences between crab cue treatments are indicated if the two corresponding bars do not share the same letter. NC = no cues, CC = chemical cues, TCC = tactile plus chemical cues.

## Discussion

This study shows that increasing dogwhelk density eliminates the limitation that a green crab can exert on dogwhelk feeding through waterborne cues released by the crab. In general, the reduction of prey feeding upon detection of predator chemical cues is thought to limit the release of waterborne metabolites by prey to reduce the attraction of predators (Barnes, 1999; Johnston et al., 2012). However, under a constant density and energetic requirements of predators, increasing prey density reduces the per-capita predation risk of prey (Ferrari et al., 2010; Guariento et al., 2015). Thus, if prey can detect conspecific density, the need for prey to reduce feeding should decrease with prey density. As dogwhelks can sense the presence of feeding conspecifics (Hughes and Dunkin, 1984; Large and Smee, 2010), the absence of crab NCEs on dogwhelk feeding at intermediate and high dogwhelk densities may therefore have resulted from dogwhelks perceiving a lower predation risk.

A separate study (Trussell et al., 2008) found that waterborne crab cues limited dogwhelk consumption of mussels despite using a higher dogwhelk density (833 dogwhelks m^-2^) than the highest density used for this study. Both that study and this one used one green crab per aquarium and fed dogwhelks to the crabs. However, the aquaria used in this study were 45 times larger than those used in the study by Trussell et al. (2008). This difference suggests that waterborne crab cues may have been more diluted in this study, in that way allowing for dogwhelk density to play a larger role and effectively limit crab NCEs. This notion is in line with studies that found that increasing predator cue concentrations often trigger stronger NCEs on prey (Loose and Dawidowicz, 1994; von Elert and Ponert, 2000; Kesavaraju et al., 2007; Ferland-Raymond et al., 2010).

The present study also clarifies the modulation of predator NCEs by prey density for the studied species assemblage. Dogwhelks feeding on barnacles slow down consumption when they detect chemical cues from green crabs, but increasing dogwhelk density does not eliminate such NCEs (Trussell et al., 2003, 2006). The results of this study support the suggestion (Trussell et al., 2008) that the higher structural complexity of mussel stands (compared with barnacle stands) may provide more refuge opportunities for dogwhelks (in addition to abundant food), facilitating the limitation of crab NCEs by dogwhelk density.

Also as predicted, crab NCEs were more influenced by dogwhelk density under crab chemical cues alone than under chemical and tactile cues combined. In fact, under chemical and tactile cues, crab NCEs always occurred regardless of dogwhelk density. This result supports the notion that prey perceive a higher predation risk when predators can physically contact the prey (albeit without consuming it) in addition to releasing waterborne cues (Luttbeg and Trussell, 2013). In such a scenario, the perceived imminence of predation risk would render prey density less relevant (or irrelevant, as found in this study) in triggering a predator avoidance response. The persistence of crab NCEs despite changes in dogwhelk density under chemical and tactile crab cues could in theory also have been influenced by the amount of chemical cues released by the mussels that were being consumed. Crabs alone consumed more mussels in the TCC environment than the dogwhelks did in the CC environment. However, dogwhelks have been found not to respond to cues from damaged mussels (Large and Smee, 2010), so the difference in such cues between the CC and TCC treatments likely had no influence.

Overall, this study reinforces the notions that prey evaluate conspecific density when assessing predation risk and that the type of cues that prey are exposed to affect their interpretation of risk. These results provide further evidence of the complexities of nonconsumptive interspecific interactions that shape aquatic communities.

## Acknowledgements

We thank the Bedford Institute of Oceanography (Dartmouth, Nova Scotia) for the logistic support to conduct this experiment. The study was funded by a research grant awarded to M. C. Wong by Fisheries and Oceans Canada (DFO) and by a Discovery grant awarded to R. A. Scrosati by the Natural Sciences and Engineering Research Council of Canada (NSERC).

